# LUHMES neurospheres as a novel model for the study of human neurotoxoplasmosis

**DOI:** 10.64898/2026.11.13.766351

**Authors:** Vanessa R. Severo, Michael J. Holmes, Ronald C. Wek, William J. Sullivan

## Abstract

*Toxoplasma gondii* is a protozoan parasite that infects animal hosts for life due to its ability to persist as latent cysts inside tissues and organs, including the brain. Current treatments fail to target tissue cysts, which can recrudesce into life-threatening acute infection, and the presence of chronic infection in the brain has been linked to neurological and behavioral anomalies. The effects of the parasite on human neurons remain poorly understood, in part because reliable experimental models are limited. Here, we describe a LUnd Human Mesencephalic (LUHMES) neurosphere model to study neuronal *T. gondii* infection for up to 21 days. We leveraged this model to compare *T. gondii* infection in developing and mature neurons and observed spontaneous differentiation of tachyzoites into bradyzoite tissue cysts within neurospheres. We further show that maturation and cell-cycle state of the neuronal population influence the developmental fate of *T. gondii*. Transcriptomic analysis revealed that responses to infection vary with neuronal developmental state, with infection-associated changes in pathways related to synaptic function, neurogenesis, immune and inflammatory responses, metabolism, homeostasis, and neurodegeneration. We also demonstrate the utility of LUHMES neurospheres as an anti-parasitic drug screening platform, showing that the small molecule salubrinal significantly reduces the number of tissue cysts. Given that LUHMES neurospheres are genetically manipulable and can be co-cultured with glial cells, this model provides for a new and versatile platform that enables studies of parasite persistence, neuron–pathogen interactions, and drug discovery.

**IMPORTANCE:** *Toxoplasma gondii* causes an incurable parasitic infection that remains in the brain for life. Studying how the parasite affects human neurons may help explain how it persists and contributes to neurological changes. We adapted a laboratory “neurosphere” model using LUHMES cells, which progress from developing to mature human dopaminergic neurons. Mirroring human brain infection, the parasite transitions into the cyst stage associated with long-term, chronic infection in the LUHMES neurospheres. By comparing changes in gene expression, we found that neuronal responses to *T. gondii* infection differ depending on their maturation state. We also show that the neurospheres can be used to screen for new drugs that target the parasite. This versatile neurosphere model provides a powerful new tool to study *T. gondii* infection in the brain.

## INTRODUCTION

*Toxoplasma gondii* is a common protozoan parasite that has achieved global distribution among warm-blooded vertebrates due to its multiple routes of transmission (1). *T. gondii* is transmitted via oocysts expelled into the environment by felid definitive hosts, tissue cysts in infected intermediate hosts, or by parasites crossing the placenta during pregnancy. Upon infection, rapidly proliferating tachyzoites convert into encysted bradyzoites, establishing a lifelong chronic infection that can reactivate during immunosuppression and become life-threatening (2). Acute infection can be controlled with anti-parasitic drugs (e.g., antifolates), but no therapy eliminates the chronic stage of *T. gondii* (3). Congenital toxoplasmosis can cause miscarriage or stillbirth, or neurodevelopmental and vision impairments in surviving newborns (4).

*T. gondii* infects any nucleated cell but preferentially forms tissue cysts in post-mitotic cells (5, 6). These cell types include neurons, retinal tissue, skeletal and cardiac muscle. The sustained presence of latent tissue cysts in the brain has led to speculation that *T. gondii* could impact neurological processes. In support of this idea, several animal species, including humans, display markedly reduced risk aversion, with rodents even losing their innate fear of felid predators (7–9). Some studies have correlated chronic toxoplasmosis with various neurological disorders (10, 11).

While *T. gondii* has been suggested to alter neuronal function through modulation of neurotransmitters and cytokines; how the infection affects human neurons remains poorly understood (12). Addressing this gap has been challenging due to the limited availability of reliable and well-defined human neuronal models for in vitro analysis. Current approaches rely predominantly on murine models, which may not truly represent the response of human neurons to infection. Some models based on human cells suffer other types of limitations. For example, human induced pluripotent stem cells (iPSCs) often generate heterogeneous cell populations that complicate the interpretation of neuron-specific responses to infection. Human cancer-derived cell lines carry the disadvantage of relatively unstable genomes and may not faithfully reproduce the physiological properties of human neurons (13, 14). Homogeneous cultures of primary human neurons typically have a limited lifespan in vitro, restricting their ability to support studies of neurotoxoplasmosis (13, 14).

To help overcome these limitations, we present a 3D human neurosphere model using LUnd Human MESencephalic (LUHMES) cells that supports *T. gondii* infection. LUHMES are human neuronal precursor cells engineered to differentiate into post-mitotic neurons following exposure to tetracycline, rapidly generating a homogeneous population of mature dopaminergic neurons (15, 16). In contrast to monolayers, 3D culture systems promote cellular self-organization that recapitulates tissue structure and function, ultimately enhancing cell survival and extending the lifespan of cells in culture. In a 3D neurosphere culture system, LUHMES cells remain viable for up to 21 days (17), greatly extending the opportunity to study the effects of human infection in vitro.

In the LUHMES neurosphere model, we observed a high degree of spontaneous tissue cyst formation upon infection, noting that the proliferative state of neuronal cells is a key determinant of *T. gondii* differentiation. We also performed a transcriptomic analysis of *T. gondii*-infected neurospheres, which revealed transcriptional changes in pathways related to synaptic function, neurogenesis, immunity, inflammation, metabolic homeostasis, and neurodegeneration. Further, we show that treatment with the eIF2α dephosphorylation inhibitor salubrinal (18, 19) reduced the tissue cyst burden in *T. gondii*-infected LUHMES neurospheres. In short, we conclude that the LUHMES neurosphere model offers a biologically relevant platform for the investigation of human neurotoxoplasmosis, host-pathogen interactions, and candidate drugs that target the chronic cyst stage.

## RESULTS AND DISCUSSION

### Infection of LUHMES neurospheres with *T. gondii*

LUHMES cells are human neuronal precursors isolated from the ventral mesencephalon that have been immortalized via transduction with a tetracycline-regulated v-myc, enabling conditional terminal differentiation into mature dopaminergic neurons (15). We initially assessed the maturation of LUHMES neuronal cells in monolayer cultures treated with 1µg/ml anhydrotetracycline (aTc) for 14 days. LUHMES began to express markers of mature neurons, βIII-tubulin and NeuN, by day 5 (Fig. S1A). LUHMES monolayers were cultured with 10 days of aTc treatment and infected with type II ME49 strain parasites. The parasite infection was monitored by IFA using anti-SAG1 and *Dolichos biflorus* agglutinin (DBA) to visualize tachyzoites and bradyzoite cysts, respectively (Fig. S1B). LUHMES monolayers supported infection and promoted spontaneous cyst formation (Fig. S1B); however, uninfected LUHMES monolayers displayed substantial decreases in viability after ∼8 days and largely died off 4 days post-infection (DPI). The short shelf life and fragility of LUHMES monolayers limit the downstream analyses that can be performed.

The lifespan of LUHMES neuronal cells can be extended to at least 3 weeks if the cells were cultured as 3D neurospheres instead of 2D monolayers (17). To recapitulate this system for *T. gondii* studies, we initiated differentiation of the LUHMES precursor cells into dopaminergic neurons by replacing the proliferation medium with differentiation medium containing aTc, a cAMP analog, and glial cell line-derived neurotrophic factor (GDNF) while being maintained on a gyratory shaker at 37°C to promote aggregation into neurospheres (Table S1). Inclusion of aTc arrests LUHMES cell proliferation by silencing v-myc expression, while the cAMP analog and GDNF promote acquisition of dopaminergic neuronal characteristics (15, 20). Taxol is added on day 3 post-differentiation (PD) and maintained for 48h to inhibit residual proliferation, promoting a homogeneous population of post-mitotic neurons (Fig. 1A, B) (17, 20). To monitor LUHMES differentiation, the expression of mature neuronal markers NF200, MAP2, βIII-tubulin, and NeuN were evaluated by IFA (Fig. S2A). In addition, we used the proliferation marker Ki-67 to mark incompletely differentiated LUHMES cells (Fig. S2B). By day 12 PD, LUHMES cells displayed markers consistent with fully mature, post-mitotic neurons, as previously described (17).

**Figure 1.**
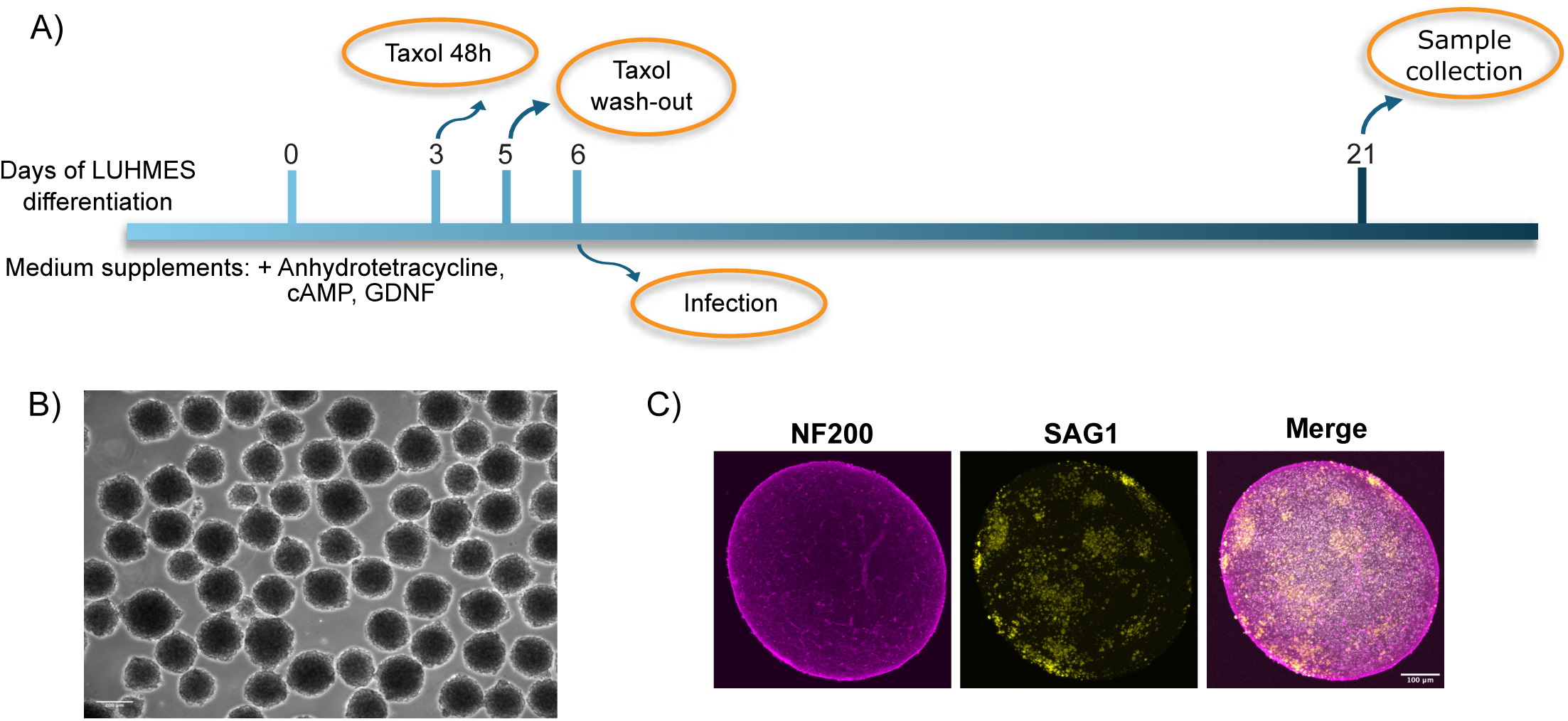
LUHMES neurospheres support *T. gondii* infection. (A) Schematic of LUHMES cell differentiation and infection. (B) Representative image of floating neurospheres following differentiation. Scale bar = 200 µm. (C) Representative immunofluorescence microscopy images of *T. gondii*-infected LUHMES neurospheres. SAG1 (yellow), NF200 (magenta), Hoechst (gray). Scale bar = 100 µm.

We initially infected LUHMES neurospheres on day 6 PD and assessed infection on 15 DPI. SAG1 staining revealed a substantial amount of tachyzoites, indicating that LUHMES neurospheres support *T. gondii* infection as previously suggested (21) (Fig. 1C).

### *T. gondii* spontaneous cyst formation occurs more frequently in post-mitotic neurons

*T. gondii* undergoes spontaneous cyst formation in post-mitotic cells that have permanently exited the cell cycle, such as mature neurons, whereas cyst formation occurs less efficiently in proliferating or quiescent cells (5, 22). We took advantage of the LUHMES model to examine *T. gondii* infection during neurosphere maturation, which includes proliferating cells, and after neuronal differentiation was complete (post-mitotic). Neurospheres infected on day 6 PD still contain Ki-67-positive cells (Ki-67^+^), which we designate as developing neurons (Fig. 2A). In contrast, neurospheres infected on day 12 PD are Ki-67-negative (Ki-67^-^), which we designate as mature neurons (Fig. 2A). This approach allows comparisons of *T. gondii* infection during neuronal development and post-neuronal development, modelling aspects of disease relevant to congenital toxoplasmosis versus adult-acquired toxoplasmosis.

**Figure 2.**
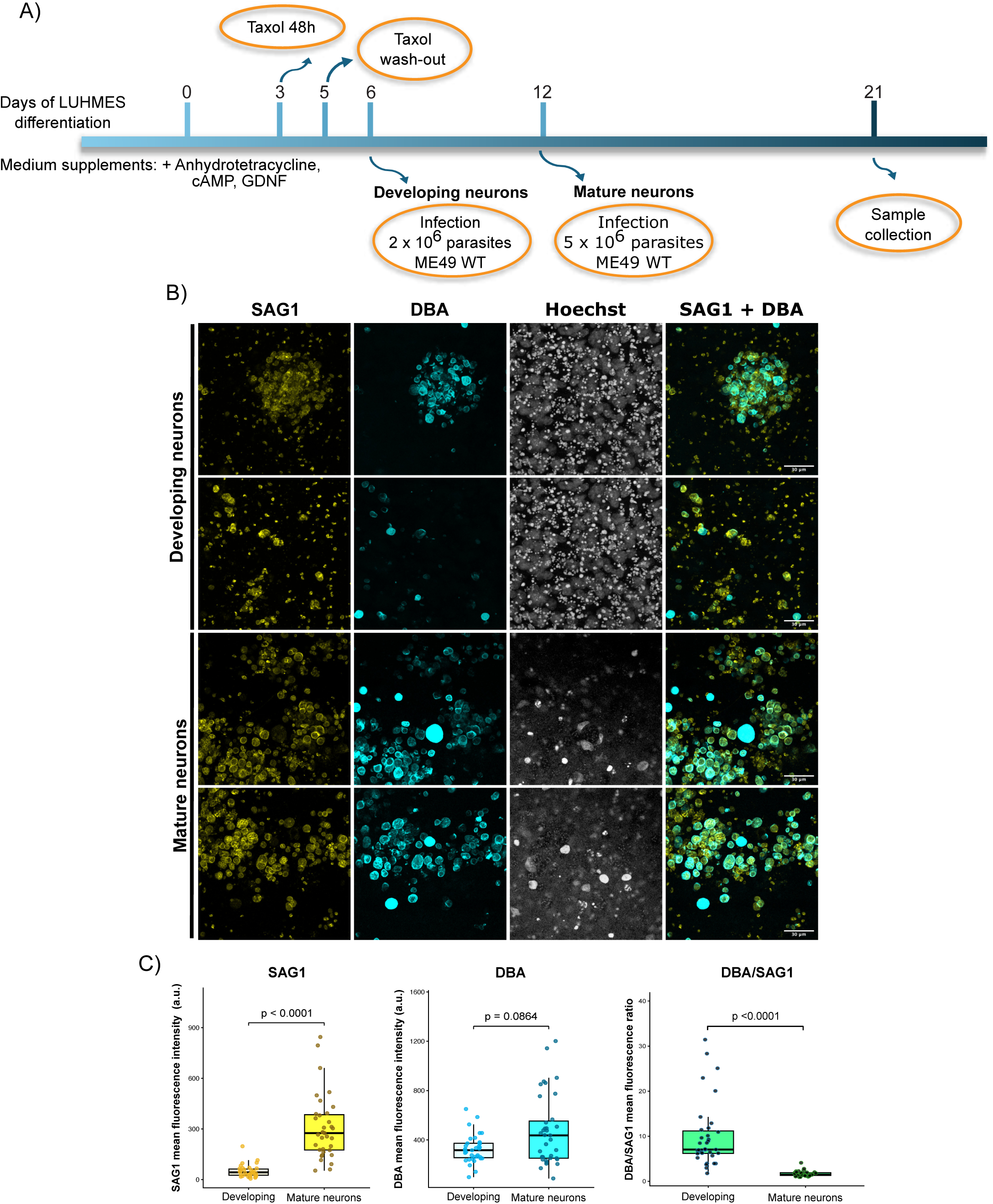
*T. gondii* infection of 3D LUHMES neurospheres leads to spontaneous cyst formation. (A) Schematic of LUHMES cell differentiation and infection. (B) Representative immunofluorescence microscopy images of infected developing and mature neurons. SAG1 (yellow), DBA (cyan), Hoechst (gray). Scale bar = 30 µm. (C) Quantification of SAG1 and DBA mean fluorescence intensity and the DBA/SAG1 mean fluorescence intensity ratio. Data are presented as boxplots, with boxes representing the interquartile range (25th-75th percentiles) and center line indicating the median. Whiskers indicate the minimum and maximum values. Statistical significance was determined using the Mann-Whitney U test.

We next examined infection of this dual model system using two different *T. gondii* strains: type I RH (largely noncystogenic) and type II ME49 (cystogenic). RH strain parasites did not form cysts in developing neurons, but some cysts formed spontaneously in mature neurons, as detected by DBA staining (Fig. S3). Appreciable SAG1 staining tachyzoites were evident in both developing and mature neurons infected with RH. However, the ME49 strain showed evidence of increasing amounts of spontaneous cyst formation over time in developing as well as mature neurons (Fig. S4A, B). Our findings are consistent with earlier reports noting that tachyzoites spontaneously convert into bradyzoites after invading post-mitotic neurons (5, 22). The remainder of our experiments employ only the ME49 strain.

In developing neurons infected with *T. gondii*, morphological changes in host nuclei consistent with cell death were observed by Hoechst staining. This observation, along with the presence of many extracellular SAG1^+^ parasites, is indicative of a more active tachyzoite infection (Fig. 2C). In contrast, parasites in mature neurons were predominantly confined to parasitophorous vacuoles or cysts, likely reflecting reduced parasite proliferation and their transition toward quiescent bradyzoites (Fig. 2C). Comparing SAG1 and DBA mean fluorescence intensity (MFI) in developing versus mature neurons revealed a significant increase in both markers, along with greater colocalization in mature neurons (Fig. 2B, C). The DBA/SAG1 MFI ratio was higher in developing neurons, potentially reflecting their longer time in culture (Fig. 2C). It should be noted that all samples were analyzed on day 21 PD, which means that developing neurons are maintained for 15 days after infection, whereas mature neurons are maintained for 9 days.

To more accurately compare infection in developing and mature neurons, we omitted the cAMP analog from the LUHMES differentiation medium to prolong the immature state. In this system, cAMP promotes neuronal maturation and dopaminergic gene expression (20). At day 12 PD, cultures without cAMP still contained proliferating cells (-cAMP, Ki-67^+^), whereas cultures with cAMP contained mature neurons (+cAMP, Ki-67^-^) (Fig S5). We infected both groups on day 12 PD with the same number of parasites (5.5×10^6^), allowing the dose of parasites and culture duration to be matched. A significant increase in DBA and concomitant decrease in SAG1 was observed in mature neurons, which led to a comparatively robust increase in the DBA/SAG1 ratio in mature neurons (Fig. 3A, B). Together, these findings affirm that maturation and cell-cycle state of the neuronal population influence the developmental fate of *T. gondii*. In agreement with our results, previous studies have demonstrated that terminally differentiated, post-mitotic myotubes support higher levels of cyst formation compared to proliferating myoblasts (5, 22).

**Figure 3.**
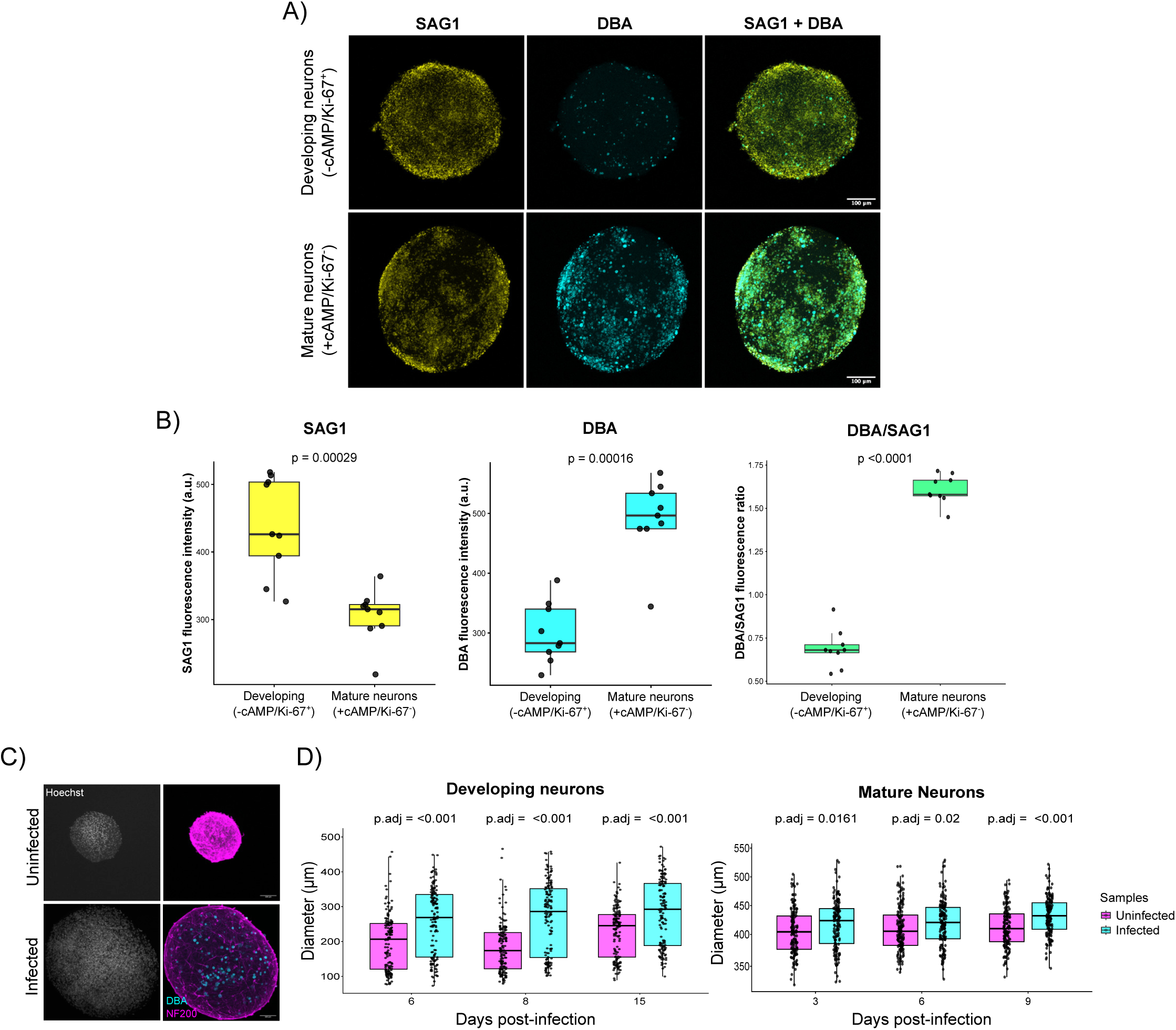
*T. gondii* forms cysts more efficiently in mature neurons and increases neurosphere size following infection. (A) Representative immunofluorescence microscopy images of neurospheres infected on day 12 PD in developing neurons (-cAMP/Ki-67^+^) and mature neurons (+cAMP/Ki-67^-^). SAG1 (yellow), DBA (cyan), Hoechst (gray). Images are max-intensity Z-projections of entire neurospheres. Scale bar = 100 µm. (B) Quantification of SAG1 and DBA mean fluorescence intensity and DBA/SAG1 mean fluorescence intensity ratio. Data are presented as mean ± standard deviation. Statistical significance was determined using the Mann-Whitney U test. (C) Representative images of infected and uninfected neurospheres visualized by immunofluorescence microscopy. Hoechst (gray), NF200 (magenta), DBA (cyan). Images are max-intensity Z-projections of entire neurospheres. Scale bar = 100 µm. (D) Neurosphere size was assessed in floating cultures using brightfield microscopy. Neurosphere diameter was measured using Fiji/ImageJ software. Data are presented as boxplots, with boxes representing the interquartile range (25th-75th percentiles) and center line indicating the median. Whiskers indicate the minimum and maximum values. Statistical significance was assessed using the Aligned Rank Transform (ART) and Wilcoxon–Mann–Whitney tests, with Holm correction for multiple comparisons.

Despite the high frequency of spontaneous cyst formation, a significant portion of SAG1^+^ parasites remain in the neurospheres. It is likely that parasite brain cyst development is augmented by factors from other cell types, such as glial cells, which can be incorporated into LUHMES neurospheres (23, 24).

### Neurospheres increase in size following *T. gondii* infection

Previous 3D infection of other cell models have reported changes in spheroid size upon *T. gondii* infection: mouse neural progenitor neurospheres and iPSCs-derived BrainSpheres became smaller after *T. gondii* infection (25, 26), as did chick skeletal muscle spheroids (27). In contrast, LUHMES neurospheres expanded in size by 6 DPI, which increased over time in developing neurons reaching 64.37% of increase by 8 DPI and 19.2% of increase by 15 DPI (Fig. 3C, D). This effect was also observed in mature neurons, although at a more modest level with 5.4% of increase by our experimental endpoints (Fig. 3D). The mechanisms underlying these observations remain unclear and warrant further investigation. As neurons depend on glial cells for survival and maintenance (28), the disparate effect infection has on these spheroid models may be associated with cellular composition. LUHMES neuropsheres are homogeneous whereas iPSCs-derived models typically contain additional cell types.

### Transcriptomic profiling of *T. gondii* infected LUHMES neurospheres

To characterize the LUHMES transcriptional response to *T. gondii* infection, we performed RNA-seq analysis of ME49-infected and uninfected LUHMES neurospheres. Samples were generated for developing and mature neurons infected with *T. gondii* following the culture regimen outlined in Fig. 2A. Uninfected samples were maintained in parallel under the same culture conditions. Principal Component Analysis (PCA) revealed clear separation between groups under both conditions, indicating distinct gene expression profiles (Fig. S6).

We obtained an average of 99.3% reads for human neurons compared to an average of 0.7% for *T. gondii* (Table 1). Despite the relatively low number of parasite transcripts detected, we were able to observe differences in stage-specific mRNAs in developing and mature neurons. Bradyzoite-associated transcripts such as BAG1, ENO1, and CST1 were expressed at higher levels in mature neurospheres; tachyzoite-associated transcripts such as SAG1 and ENO2 were more abundant in developing neurospheres (Fig. 4A). These transcriptome changes largely align with the DBA/SAG1 ratio on Fig. 3B, showing a greater frequency of spontaneous cyst formation in mature neurons compared to developing neurons.

**Figure 4.**
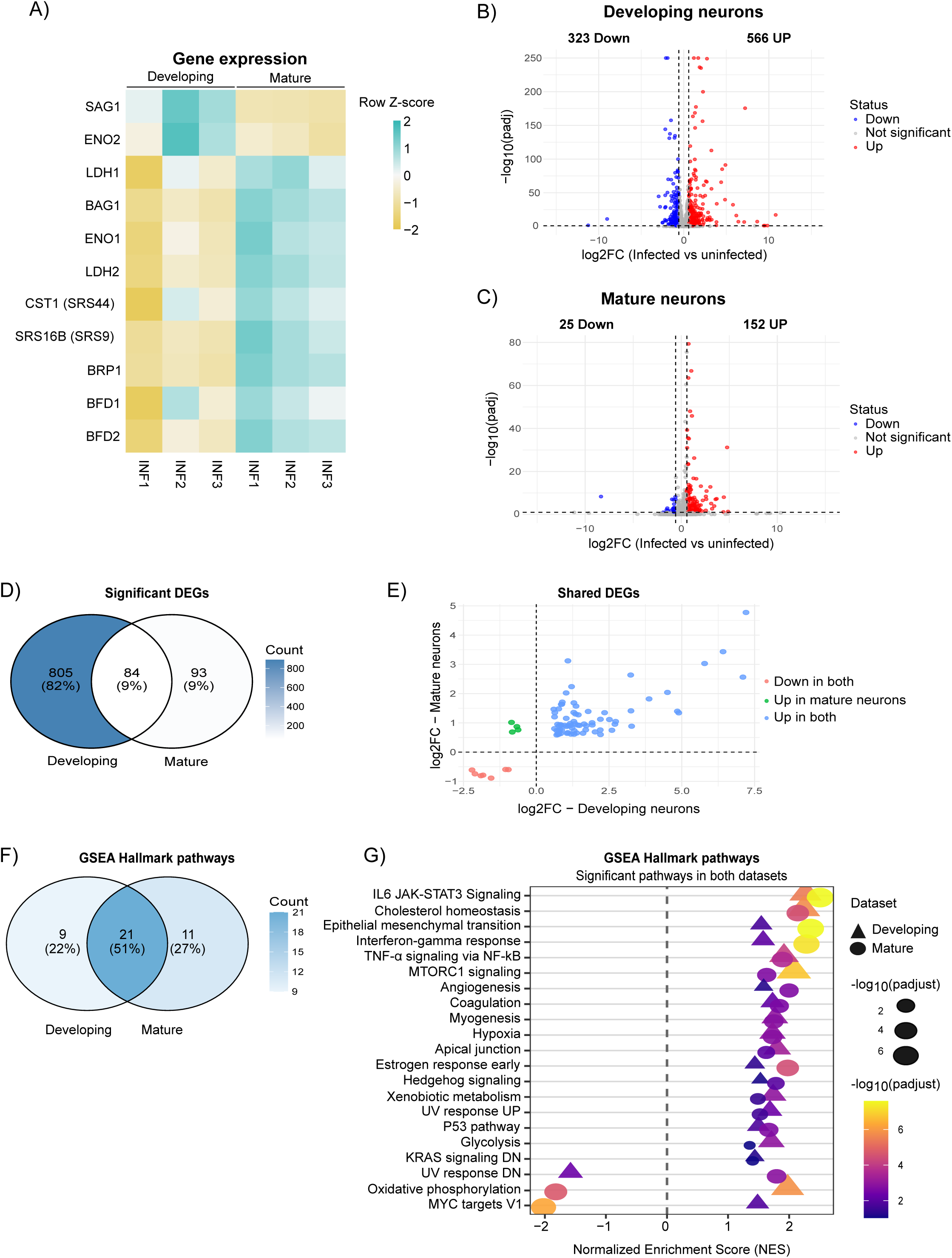
RNAseq analysis reveals *T. gondii* stage-specific gene expression and limited overlap in transcriptional changes between infected developing and mature neurons. (A) Heatmap showing gene expression across three independent experiments for developing and mature neurospheres infected with *T. gondii*. Transcript abundance is expressed as transcripts per million (TPM), transformed as log2(TPM+1), and standardized independently for each gene by calculating row Z-scores across all samples. Colors represent relative expression levels for each gene, ranging from lower relative expression (yellow) through the gene mean (white) to higher relative expression (cyan). Genes and samples are displayed without hierarchical clustering. (B) Volcano plot of differentially expressed genes (DEGs) in infected versus uninfected samples in developing and (C) mature neurons. (D) Venn diagram comparing significant DEGs between the two neuronal maturation stages. (E) Comparison of log2 fold-change (log2FC) values for DEGs shared between developing and mature neurons. (F) Venn diagram showing the overlap of significantly enriched Hallmark pathways between developing and mature neurons. (G) Plot of normalized enrichment scores (NES) and - log10(padjust) values for the Hallmark pathways.

**Table 1.** Number of reads assigned to *Homo sapiens* or *Toxoplasma gondii* genes in developing and mature infected neurons.

|  | **Developing neurons** |  | **Mature neurons** |  |
|---|---|---|---|---|
| **Sample** | Read counts estimated by Salmon | Percentage | Read counts estimated by Salmon | Percentage |
| ***Homo sapiens*** |  |  |  |  |
| **INF1** | 32498653.46 | 99.64 | 34040838.7 | 99.06 |
| **INF2** | 323227049.34 | 99.18 | 34588321.2 | 99.20 |
| **INF3** | 33440318.72 | 99.42 | 29589799.6 | 99.29 |
| ***Toxoplasma gondii*** |  |  |  |  |
| **INF1** | 117627.44 | 0.36 | 320005.1 | 0.93 |
| **INF2** | 267602.51 | 0.82 | 276404.9 | 0.79 |
| **INF3** | 193879.32 | 0.57 | 211325.3 | 0.71 |

Regarding the host transcriptome changes in response to infection, we found that 566 genes were upregulated and 323 downregulated in developing neurons (Fig. 4B). There were reduced levels of gene expression changes by infection of mature neurospheres, with 152 genes upregulated and 25 downregulated (Fig. 4C). Developing and mature neurospheres infected with *T. gondii* shared 84 differentially expressed genes (DEGs), 73 of which were upregulated in both conditions (Fig. 4D, E).

The enhanced transcriptional response in developing neurons may be reflective of the more active infection compared to greater parasite quiescence as cysts in mature neurospheres. In other words, as more tachyzoites convert into latent bradyzoites, changes in host cell gene expression largely abate. The transcriptional alterations that are maintained in mature neurospheres suggest that parasite persistence continues to influence neuronal homeostasis despite the reduced magnitude of the host transcriptional response. It is important to note that parasite transcripts counts are similar in developing and mature neurons, indicating a comparable overall parasite burden in both conditions (Table 1).

Gene Set Enrichment Analysis (GSEA) showed 21 (51%) Hallmark pathways that overlap between developing versus mature neurons infected with *T. gondii* (Fig. 4F). These include pathways known to be manipulated by *T. gondii*, such as IL-6/JAK/STAT3 signaling, cholesterol homeostasis, the IFN-γ response, TNF-α signaling via NF-kB, mTORC1 signaling, glycolysis, oxidative phosphorylation (Fig. 4G) (29–33).

GSEA of biological processes in developing and mature neurospheres infected with *T. gondii* identified 28% of the processes shared by both conditions were related to synaptic function and neurogenesis, immunity and inflammation, metabolism, and homeostasis (Fig. 5A). Many of these processes were positively enriched in both conditions (Table S3/S4). Shared biological processes related to neurodevelopment and neuronal differentiation, cognition, and neuronal function showed opposite enrichment directions within conditions (Fig. 5B). They were negatively enriched in infected developing neurons and positively enriched in infected mature neurons (Fig. 5B). These findings are consistent with experimental evidence that congenital *T. gondii* infection during fetal brain development disrupt neurodevelopment and neurogenic processes, potentially contributing to central nervous system abnormalities (26, 34).

**Figure 5.**
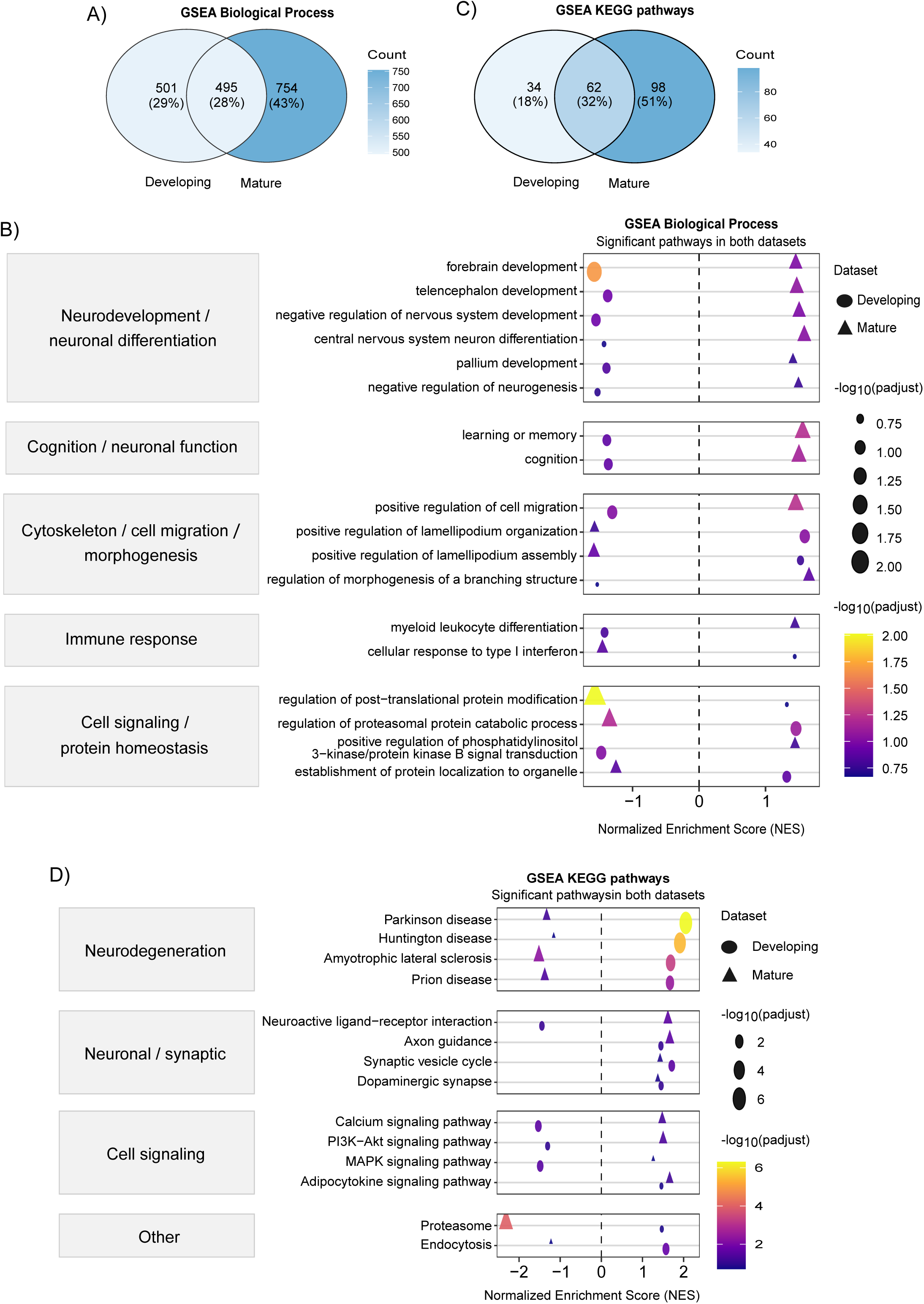
Gene Set Enrichment Analysis (GSEA) of RNAseq data identified biological processes and KEGG pathways shared between developing and mature neurons. (A) Venn diagram showing the overlap of significantly enriched biological processes between infected developing and mature neurosheres. (B) Plot of normalized enrichment scores (NES) and - log10(padjust) values for the biological process pathways. (C) Venn diagram showing the overlap of significantly enriched KEGG pathways between infected developing and mature neurospheres. (D) Plot of normalized enrichment scores (NES) and −log10(padjust) values for the KEGG pathways.

GSEA for KEGG pathways analysis identified a shared response to infection in developing and mature neurons, including changes in pathways associated with neurodegenerative diseases (Fig. 5C, D). However, these pathways were positively enriched in infected developing neurons and negatively enriched in infected mature neurons, suggesting that *T. gondii* infection may broadly perturb pathways associated with neurodegenerative disorders. (Fig. 5D; Table S3/S4). This result agrees with prior studies that showed that *T. gondii* infection alters neurodegeneration-associated pathways in primary human stem cells (35).

Overall, our transcriptomic analysis indicates that human neuronal responses to *T. gondii* infection are influenced by their developmental state. Moreover, *T. gondii* infection induces host transcriptional changes in pathways associated with synaptic function, neurogenesis, immune and inflammatory responses, metabolism, homeostasis, and neurodegeneration. To our knowledge, these datasets represent the first transcriptomic analysis of *T. gondii* infection in a pure culture of human neurons.

### LUHMES neurosphere culture as a platform for anti-toxoplasmosis drug screening

There are no approved therapies capable of eliminating latent tissue cysts, which constitute the primary obstacle to toxoplasmosis treatment. We tested whether the LUHMES neurosphere model could provide a new tool for drug screening aimed at identifying lead molecules that reduce *T. gondii* cysts in human neural cells. LUHMES cells are widely used as an in vitro model system for neurotoxicity testing and high-throughput drug screening, particularly for Parkinson’s disease (36, 37).

We previously reported in *T. gondii* infected human foreskin fibroblasts (HFFs) that an inhibitor of TgIF2**α** dephosphorylation, salubrinal, exerts activity against both tachyzoites and bradyzoites (18, 19). To evaluate the effect of salubrinal in the LUHMES neurosphere model, we treated both developing and mature neurons infected with ME49 strain *T. gondii.* In developing neurons, treatment was initiated 5 DPI with 8 µM salubrinal, the concentration previously shown to be lethal to *T. gondii* in HFFs (18). The culture medium was replaced every other day with fresh salubrinal, and samples were collected 5 or 10 days post-treatment (DPT) for IFA analysis using anti-SAG1 and DBA (Fig. 6A). We saw nearly 20% DBA reduction representing bradyzoites, but no change was observed in SAG1-marked tachyzoites in the infected developing neurons (Fig. 6B).

**Figure 6.**
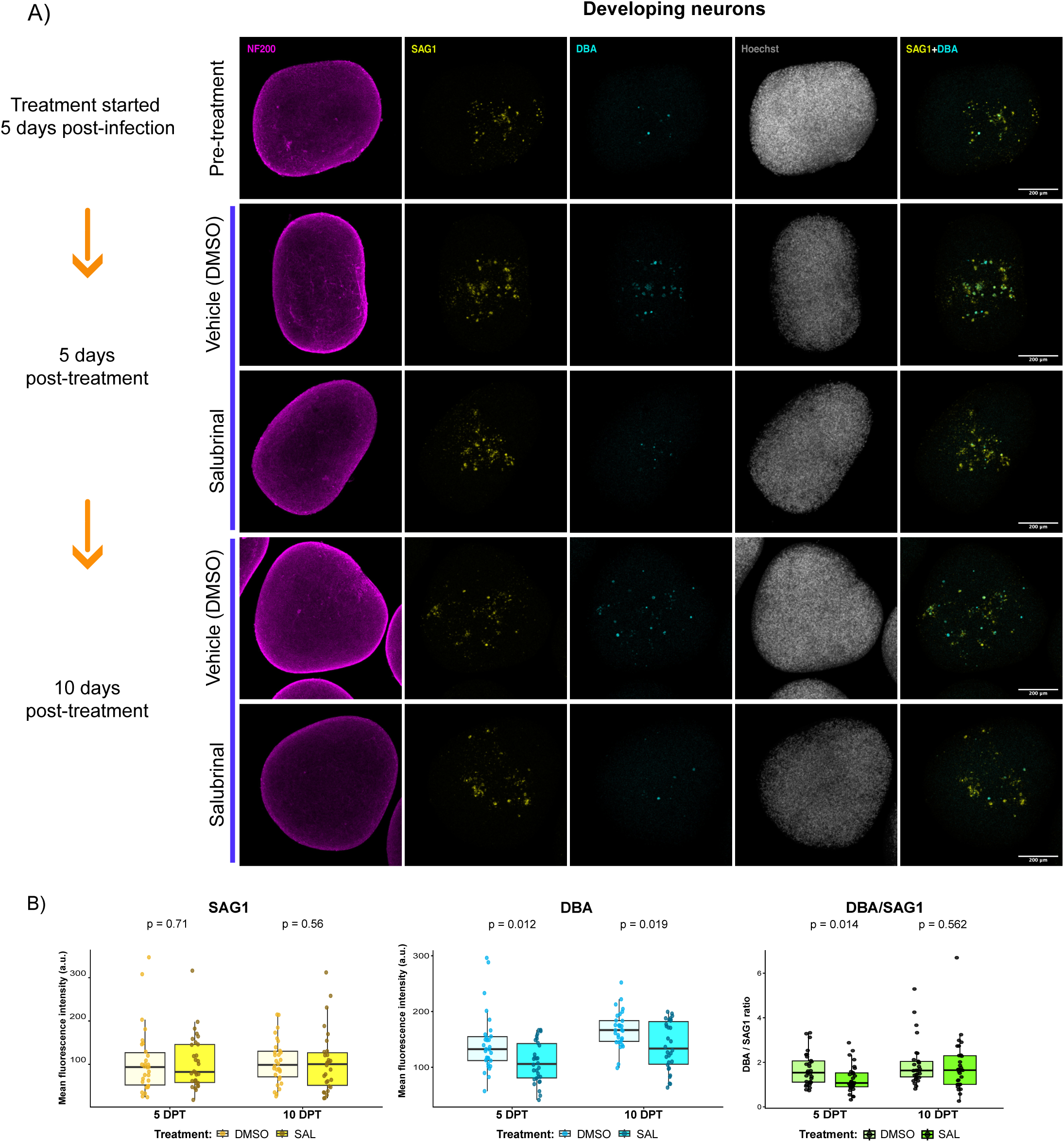
Salubrinal treatment reduces cyst burden in developing neurospheres. (A) Treatment schematic and representative immunofluorescence microscopy images acquired before treatment and at 5 and 10 days after treatment with 8 µM salubrinal or vehicle control (DMSO). NF200 (magenta), SAG1 (yellow), DBA (cyan), Hoechst (gray). Images are max-intensity Z-projections of entire neurospheres. Scale bar = 200 µm. (B) Quantification of SAG1 and DBA mean fluorescence intensity and DBA/SAG1 mean fluorescence intensity ratio of salubrinal- and DMSO-treated samples at 5 and 10 days post-treatment (DPT). Data are presented as boxplots, with boxes representing the interquartile range (25th-75th percentiles) and center line indicating the median. Whiskers indicate the minimum and maximum values. Statistical significance was assessed using the Aligned Rank Transform (ART) and Wilcoxon–Mann–Whitney tests, with Holm correction for multiple comparisons.

We also evaluated the effects of salubrinal on mature neurons infected with *T. gondii*. After 4 days of infection, the mature neurosphres were treated with 8 µM salubrinal for 5 days (Fig. 7A). As in developing neurons, we observed a nearly 61% reduction of cysts in mature neurons 5 DPT (Fig. 7B). In contrast to developing neurons, SAG1 was reduced 43% in mature neurons (Fig. 7B). This result could be reflective of the higher frequency of cyst conversion in mature neurons; alternatively, salubrinal may affect tachyzoites differently depending on neuronal maturity. Through visual assessment, we detected no overt adverse effects of salubrinal on the viability of the neurospheres, which is another attractive feature for drug screening. These findings support the utility of LUHMES neurospheres as a platform for screening anti-*T. gondii* drug candidates. The data also affirm the promise of salubrinal as a cyst-reducing agent.

**Figure 7.**
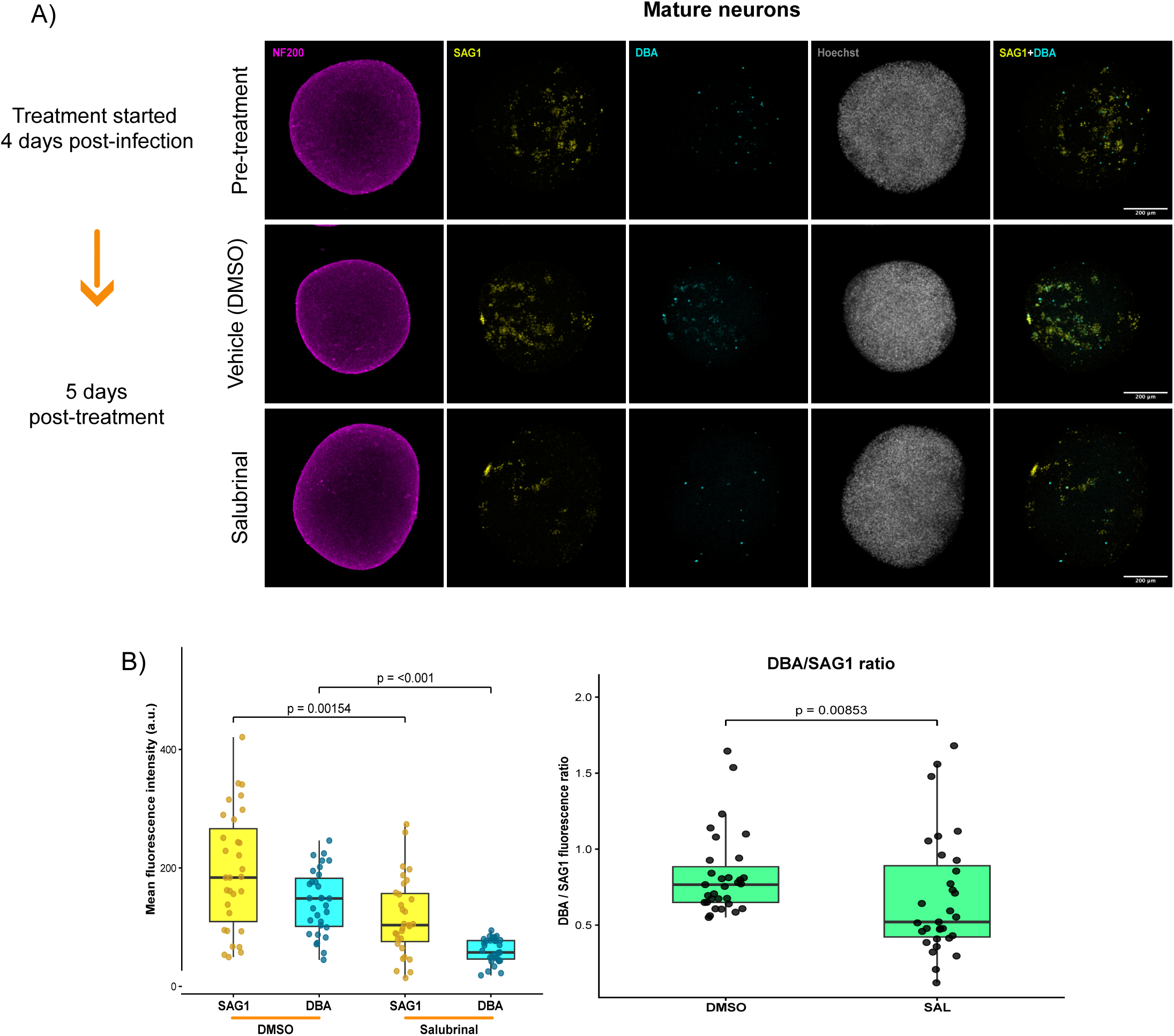
Salubrinal treatment reduces tachyzoites and cyst burden in mature neurospheres. (A) Treatment schematic and representative immunofluorescence microscopy images acquired before treatment and after 5 days of treatment with 8 µM salubrinal or vehicle control (DMSO). NF200 (magenta), SAG1 (yellow), DBA (cyan), Hoechst (gray). Images are max-intensity Z-projections of entire neurospheres. Scale bar = 200 µm. (B) Quantification of SAG1 and DBA mean fluorescence intensity and DBA/SAG1 mean fluorescence intensity ratio in salubrinal- and DMSO-treated samples. Data are presented as boxplots, with boxes representing the interquartile range (25th-75th percentiles) and center line indicating the median. Whiskers indicate the minimum and maximum values. Statistical significance was determined using the Mann-Whitney U test.

### Concluding remarks

This study establishes LUHMES neurosphere as a novel model for investigating *T. gondii* infection in human neurons. We demonstrate that this model can be used to study *T. gondii* effects on either developing or mature neurons for at least 15 to 9 days, respectively. As reported for other post-mitotic cells, a portion of tachyzoites spontaneously differentiate into latent tissue cysts after invading neurospheres, opening the door to interrogating this clinically relevant life cycle stage in the context of human neurotoxoplasmosis. Importantly, this model is amenable to screening for compounds that exert activity against tachyzoites and bradyzoites infecting human neurons. The rapid and homogeneous differentiation of LUHMES cells into dopaminergic neurons, together with their commercial availability, reproducibility, compatibility with genetic manipulation, and potential for co-culture with other cell types, makes this system an accessible and versatile tool for the *T. gondii* research community. Our initial studies show that LUHMES neurospheres hold great promise to reveal new insights into *T. gondii* persistence, neuron-pathogen interactions, and neurological effects caused by infection.

## MATERIALS AND METHODS

### LUHMES proliferation

Media compositions for LUHMES cultures are shown in Table S1. For LUHMES cell propagation, Nunc™ EasYFlask™, Nunclon Delta Surface Cell Culture Flasks (ThermoFisher Scientific) were pre-coated with a solution containing 50 µg/mL Poly-L-Ornithine (PLO) and 1 µg/mL fibronectin in Milli-Q H_2_O and incubated overnight at 37°C. Flasks were then washed twice with sterile distilled water and dried under a laminar flow bench before use. LUHMES cells (ATCC CRL-2927) were maintained in proliferation medium as previously described (38, 39).

### LUHMES differentiation and neurosphere culture

#### 2D culture

For LUHMES differentiation in 2D culture, cells were seeded onto glass coverslips pre-coated as described above. Proliferation medium was replaced by differentiation medium containing aTc and GDNF, and cultures were kept in 5% CO_2_ at 37°C. Differentiated neurons were infected on day 10 PD.

#### 3D culture

For neuronal differentiation and neurospheres formation, cells were trypsinized using TripleE Express (Life Technologies) upon reaching 70-80% confluency. A single-cell suspension was prepared in differentiation medium, and cells were seeded in 6-well plates at a density of 5.5 x10^5^ cells per well in 2 mL of differentiation medium. Plates were incubated at 37°C, 10% CO_2_ while shaking at 80 rpm on a gyratory shaker. On day 3 PD, neurospheres were treated with 10 nM Taxol (Sigma) for 48h. Neurospheres were then washed to remove Taxol before adding differentiation medium. Thereafter, differentiation medium was replenished every other day by removing 1 mL from each well and adding 1.2 mL of fresh medium. For *T. gondii* infection, 2 x 10^6^ parasites were added to each well on day 6, or 5 x 10^6^ parasites on day 12 PD. To facilitate infection, the neurospheres were incubated without shaking, then returned to the shaker after 1h.

### Parasite cultures

Wild-type ME49 and RHΔKu80 *T. gondii* parasites were provided by Dr. Vern Carruthers (University of Michigan Medical School) and maintained in Human Foreskin Fibroblasts cells (HFFs – ATCC SCRC-1041) in Dulbecco’s modified Eagle medium (DMEM; pH 7,4) supplemented with 10% fetal bovine serum (FBS), 100 U/mL penicillin, 100 µg/mL streptomycin. Infected HFF monolayers were cultured at 37°C and 5% CO_2_.

### Salubrinal treatment

Salubrinal (Tocris Bioscience Cat #2347) treatment was initiated 4 or 5 days post-infection, as indicated. Salubrinal was dissolved in DMSO and applied at a final concentration of 8 µM salubrinal in LUHMES differentiation medium. The drug and medium was refreshed every other day. Vehicle-treated samples were included as controls.

### Immunofluorescence and imaging

Immunofluorescence assays (IFA) of LUHMES neurospheres were performed as previously described (38, 39). Briefly, neurospheres were transferred to a 24-well plate, washed once in 1X PBS, and fixed for 1h in 4% paraformaldehyde. Neurosphere cultures were then washed twice and incubated in blocking solution for 1h at 4°C, followed by incubation with primary antibodies diluted in blocking solution for 48h at 4°C on a shaker (solutions and antibodies information are listed in Table S1). Samples were washed 3 times in wash solution I before incubating for 24h at 4°C with secondary antibodies and dyes diluted in 1X PBS. Samples were then washed twice in wash solution II, and once in 1X PBS. Slides were mounted with Immu-Mount (Epredia #9990402) and viewed using Zeiss LSM 800 AxioObserver microscope with an Airyscan detector. The Airyscan processing was performed using Zen Blue (version 2.6.76; Zeiss, Oberkochen, Germany). Mean Fluorescence Intensity (MFI) of SAG1 and DBA was quantified using QuPath software (version 0.7) from 15 random fields across at least two independent experiments (40). The ratios were calculated by dividing DBA MFI by SAG1 MFI values. For neurosphere size measurement, brightfield images of floating neurospheres were acquired using a Leica DMI6000B microscope using a 5X objective. Diameters of 50 neurospheres from 3 independent experiments were measured using Fiji/ImageJ software (41).

### RNA-seq and data analysis

RNA-seq was conducted using three biological replicates per condition. Neurospheres were infected on day 6 or 12 PD and collected on day 21 PD (Fig. 2A). Samples were washed twice with 1X PBS and resuspended in TRIzol reagent (Invitrogen) according to the manufacturer’s protocol. Library preparations were carried out by the Center for Medical Genomics at Indiana University School of Medicine using standard Illumina protocols. Raw RNA-seq reads in FASTQ format were initially subjected to quality assessment using FastQC, and the resulting reports were aggregated and summarized with MultiQC to evaluate sequencing quality metrics across all samples. Adapter sequences and low-quality bases were removed using fastp with default parameters, generating high-quality reads for downstream analyses (42). Transcript quantification was performed using Salmon (version 1.11.4) in quasi-mapping mode (43). A Salmon index was built from a concatenated human-*T.gondii* reference transcriptome comprising GENCODE release 49, and the ToxoDB release 68 *T. gondii* annotated transcripts (44, 45). Cleaned reads were mapped to the indexed concatenated transcriptome, and transcript abundances were estimated as transcript-level counts and transcripts per million (TPM). Transcript-level quantifications were imported into R (version 4.5.3) (46, 47) and summarized to gene-level counts using the Bioconductor package tximeta (version 1.28.3) (48). Differential gene expression analysis was subsequently performed using the DESeq2 package (version 1.50.2) (49). Genes exhibiting significant expression changes were identified based on adjusted *p*-values corrected for multiple testing using the Benjamini–Hochberg method, and genes were considered differentially expressed when they met the thresholds of adjusted *p*-value < 0.1 and absolute log2 fold change (|log2FC|) > 0.585, corresponding to a minimum 1.5-fold change in gene expression (Fig. S4/S5). For Gene Set Enrichment Analysis (GSEA), clusterProfiler package with default parameters (version 4.18.4) was used (50). Data visualization, including principal component analysis (PCA), volcano plots, and other graphical representations, was performed in R using the ggplot2 package (4.0.3) and associated visualization tools (51). Sequencing libraries were read at 2 x 150 bp and can be accessed at GSE353347.

## ACKNOWLEDGMENTS

The authors thank Dr. Anita Koshy for helpful discussions and Rian DelVecchio for technical assistance. This research was supported by a grant from the National Institutes of Health (AI172752 to WJS and RCW). Additionally, support was provided by the Indiana Clinical and Translational Sciences Institute funded, in part by Grant Number UM1TR004402 from the National Institutes of Health, National Center for Advancing Translational Sciences, Clinical and Translational Sciences Award (to VRS). The content is solely the responsibility of the authors and does not necessarily represent the official views of the National Institutes of Health.

## SUPPLEMENTAL TABLE LEGENDS

**Table S1. List of reagents**

**Table S2. DESeq2**

**Table S3. GSEA - Developing neurons**

**Table S4. GSEA - Mature neurons**

## SUPPLEMENTAL FIGURE LEGENDS

**Figure S1. LUHMES differentiation and *T. gondii* infection in 2D culture.** (A) Representative immunofluorescence microscopy images of LUHMES cells at different days of differentiation. The neuronal maturation markers βIII-tubulin (magenta) and NeuN (yellow) were used to assess differentiation. Nuclei were stained with Hoechst (blue). Phase-contrast images are shown. Scale bar = 20 µm. (B) Representative immunofluorescence microscopy images of 2D LUHMES culture at different days post-infection (DPI). F-actin was labeled with phalloidin (magenta). SAG1 (yellow), DBA (cyan), Hoechst (blue). Scale bar = 20 µm.

**Figure S2. Complete neurosphere maturation is achieved by 12 days post-differentiation.** (A) Representative immunofluorescence microscopy images showing the expression of mature neuronal markers. MAP2, βIII-tubulin, and NF200 are shown in magenta; NeuN is shown in yellow; and nuclei are stained with Hoechst in grey. (B) Representative immunofluorescence microscopy images of neuronal maturation at 6, 12, 15, and 21 days PD. The proliferation marker Ki-67 (yellow) was used to assess cell-cycle exit and neuronal maturation was evaluated using NF200 (magenta). Nuclei stained with Hoechst are shown in gray. Images are max-intensity Z-projections of entire neurospheres. Scale bar = 150 µm.

**Figure S3. The RH strain forms cysts in mature neurons but not in developing neurons.** Representative immunofluorescence microscopy images of developing and mature neurons infected with the RH strain. SAG1 (yellow), DBA (cyan). Top panel images are max-intensity Z-projections of entire neurospheres. Scale bar sizes are indicated on the micrographs.

**Figure S4. Course of infection in developing and mature neurons.** Representative immunofluorescence microscopy images at different days post-infection in (A) developing and (B) mature neurons. NF200 (magenta), SAG1 (yellow), DBA (cyan), Hoechst (gray). Images are max-intensity Z-projections of entire neurospheres. Scale bars: 100 µm for developing neurons and 200 µm for mature neurons.

**Figure S5. Removing the cAMP analog can delay neurosphere maturation.** Representative immunofluorescence microscopy images of neuronal maturation at 6- and 12-days PD with or without cAMP. The proliferation marker Ki-67 (yellow) was used to assess cell-cycle exit and neuronal maturation was evaluated using NF200 (magenta). Nuclei stained with Hoechst are shown in gray. Images are max-intensity Z-projections of entire neurospheres. Scale bar = 200 µm.

**Figure S6. RNAseq principal component analysis (PCA).** (A) RNAseq PCA plot of infected versus uninfected developing neurospheres. (B) RNAseq PCA plot of infected versus uninfected mature neurospheres.

